# Dual roles of crustacean female sex hormone during juvenile stage in the kuruma prawn *Marsupenaeus japonicus*

**DOI:** 10.1101/2023.05.13.540635

**Authors:** Kenji Toyota, Hanako Matsushima, Rei Osanai, Tomoyuki Okutsu, Fumihiro Yamane, Tsuyoshi Ohira

**Author notes:** Correspondences: Kenji Toyota, Noto Marine Laboratory, Institute of Nature and Environmental Technology, Kanazawa University, Ogi, Noto-cho, Ishikawa 927-0553, Japan., Prof. Tsuyoshi Ohira, Department of Biological Sciences, Faculty of Science, Kanagawa University, 2946 Tsuchiya, Hiratsuka, Kanagawa, 259-1293, Japan.

## Abstract

The crustacean female sex hormone (CFSH) has been identified as a female-specific hormone that plays a crucial role in female phenotype developments in the blue crab *Callinectes sapidus*. To date, its homologous genes have been reported in various decapod species. Additionally, unlike the blue crab, several species have two different CFSH subtypes. The kuruma prawn *Marsupenaeus japonicus* is a representative example species of this phenomenon, having two CFSH subtypes identified from the eyestalk (MajCFSH) and ovary (MajCFSH-ov). Eyestalk-type MajCFSH is expressed predominantly in the eyestalk at the same level in both sexes, indicating no female-specificity. Here, we conducted gene knockdown analysis of eyestalk-type MajCFSH using sexually immature juveniles of kuruma prawn (average body length: ∼10 mm) to elucidate its physiological functions. As a result, MajCFSH-knockdown did not affect the development of sex-specific characteristics such as external reproductive organs, while it induced apparent growth suppression in male juveniles, implying that MajCFSH may play a male-specific juvenile growth role. Moreover, MajCFSH-knockdown female and male juveniles changed their body color to become brighter, indicating that MajCFSH has the ability to change body color by dispersing the pigment granules in the chromatophore. Overall, our present study improved our understanding of the physiological roles of CFSH using kuruma prawn.

**Highlights:**

- MajCFSH-knockdown was conducted using sexually immature juveniles of kuruma prawn.
- MajCFSH-knockdown male juveniles showed significant growth suppression.
- MajCFSH-knockdown female and male juveniles changed their body color to become brighter.

## Introduction

The kuruma prawn (*Marsupenaeus japonicus*) is widely distributed in the Indo-West Pacific region (Holthuis, 1980). Sexually-mature kuruma prawns spawn offshore and, after hatching, the post-larvae migrate toward inner bays and settle on tidal flats after a planktonic period. Post-larvae have little ability to submerge in sand, so they stay near the shoreline to escape predator attacks, and move around the tidal flats according to the tides. After growing above a certain size on the tidal flats, the juvenile prawns move to deeper waters offshore to mature sexually. Thus, depending on their growth stage, kuruma prawns use a wide range of seawaters as their habitat, from tidal flat areas in inner bays to offshore areas (Sakaji and Nishimoto, 2022).

Despite the economic importance of this species, its annual catches have declined sharply since the 1990s (Hamasaki and Kitada, 2013). To change this situation, various researches about seed production and aquaculture of the kuruma prawn were conducted (Hamasaki and Kitada, 2006). The establishment of techniques for rearing juvenile kuruma prawns and the ability to produce seedlings on a large scale has led to the development of aquaculture and cultivated fisheries both domestically and abroad (Hamasaki and Kitada, 2013). Since the demand for important fishery species including kuruma prawn is expected to increase worldwide in the future, there is a need to develop new technologies that will make aquaculture augmentation technologies even more efficient. To achieve this goal, several studies have been conducted to analyze the transcriptome (Zhong et al., 2017) and genome assemblies of kuruma prawn (Kawato et al., 2021), and decapod crustaceans have been widely studied to develop sex control techniques and conduct research on growth promotion (Levy and Sagi, 2020).

The crustacean female sex hormone (CFSH) has been identified as a female-specific hormone that plays a crucial role in female phenotype developments in the blue crab *Callinectes sapidus* (Zmora and Chung, 2014). Gene knockdown of CFSH resulted in the anomalous development of female reproductive characteristics such as ovigerous setae and gonopores in *C. sapidus* males (Zmora and Chung 2014), and the mud crab *Scylla paramamosain* (Jiang et al. 2020). To date, its homologous genes have been reported in various decapod species such as the green shore crab, *Carcinus maenas* (Oliphant et al. 2018), the crayfish *Procambarus clarkii* (Veenstra 2015), the peppermint shrimp *Lysmata vittata* (Liu et al. 2021), the banana shrimp *Fenneropenaeus merguiensis* (Powell et al. 2015), and the giant river prawn *Macrobrachium rosenbergii* (Suwansa-Ard et al. 2015). Additionally, unlike the blue crab, several species have two different CFSH subtypes. The kuruma prawn is a representative example species of this phenomenon, having two CFSH subtypes identified from the eyestalk (MajCFSH) and ovary (MajCFSH-ov) (Kotaka and Ohira 2018; Tsutsui et al. 2018). These two MajCFSH subtypes differ not only in their primary gene structures, but also in their expression sites. Eyestalk-type MajCFSH is expressed predominantly in the eyestalk, however, it is expressed at the same level in both sexes, indicating no female-specificity (Kotaka and Ohira 2018). Ovary-type Maj-CFSH-ov is predominantly expressed in oogonia and previtellogenic oocytes during vitellogenesis, indicating that it may take part in reproductive processes (Tsutsui et al. 2018). However, the differences in physiological function between MajCFSH and MajCFSH-ov subtypes remain unclear. Here, we conducted gene knockdown analysis of eyestalk-type MajCFSH using sexually immature juveniles of kuruma prawn (average body length: ∼10 mm) to elucidate its physiological functions.

## Materials and methods

### Ethical statement

The following experiments were performed according to the Guidelines for the Care and Use of Experimental Animals of Kanagawa University by the Animal Care and Use Committee of Kanagawa University, where they were conducted. No specific permissions are required for studies that do not involve endangered or protected invertebrate species. All efforts were made to minimize the suffering of the animals.

### Animals

The kuruma prawn (*Marsupenaeus japonicus*) used in this study were produced at Aichi Prefectural Sea Farming Center (Aichi, Japan) in July 2021. Juvenile prawns were transferred to Mie Prefectural Fish Farming Center (Mie, Japan) and experiments were conducted there. Eighty mixed-sex prawns of each injection group were kept in a 100 L black tank with natural seawater (23-27°C) under natural daylight. The commercial prawn diet, Vitalprawn (Higashimaru Co., Ltd., Kagoshima, Japan), was fed until the end of the experiments.

### RNA interference and measurement of several body traits

Knockdown of MajCFSH was performed using the RNA interference (RNAi) technique. The sequence of MajCFSH was retrieved from the NCBI database (accession: LC224021). The green fluorescence protein (GFP) gene was used as a negative control. Double-stranded RNA (dsRNA) was synthesized using the MEGAscript RNAi Kit (Life Technologies, Carlsbad, CA, USA) **(Table 1)**. A total of 80 juveniles at 17 days after larval metamorphosis (post-larvae 17: PL17) with an average body length of about 10 mm, which could not be distinguished as female or male by external morphology (theoretical sexual ratio, 50:50), were used in both the MajCFSH and GFP groups. The dsRNA was injected into the body cavity of juveniles at a concentration of 0.5 μg/10 μl/individual once every other week for six weeks (injections: 1, 3, 5 weeks) using an ultrafine needle (ITO CORPORATION, Shizuoka, Japan). After six weeks, the body length and body weight were measured, and sexual identity was observed based on abdominal morphology using all surviving juveniles (GFP-dsRNA: 33 juveniles, MajCFSH: 44 juveniles). Photographs were then taken from the dorsal side for body color measurement (TG-6, Olympus, Tokyo, Japan). Three sites (P1-P3, Figure S1A) were defined as the points for extraction of color information, and these colors were transformed into the RGB (Red, Green, Blue) color space. Then, average RGB values were transformed into HSV (Hue, Saturation, Value) color space using Photoshop (Adobe KK, Tokyo, Japan). Visualization and statistics were conducted using R (R Core Team, 2022).

**Table 1.**
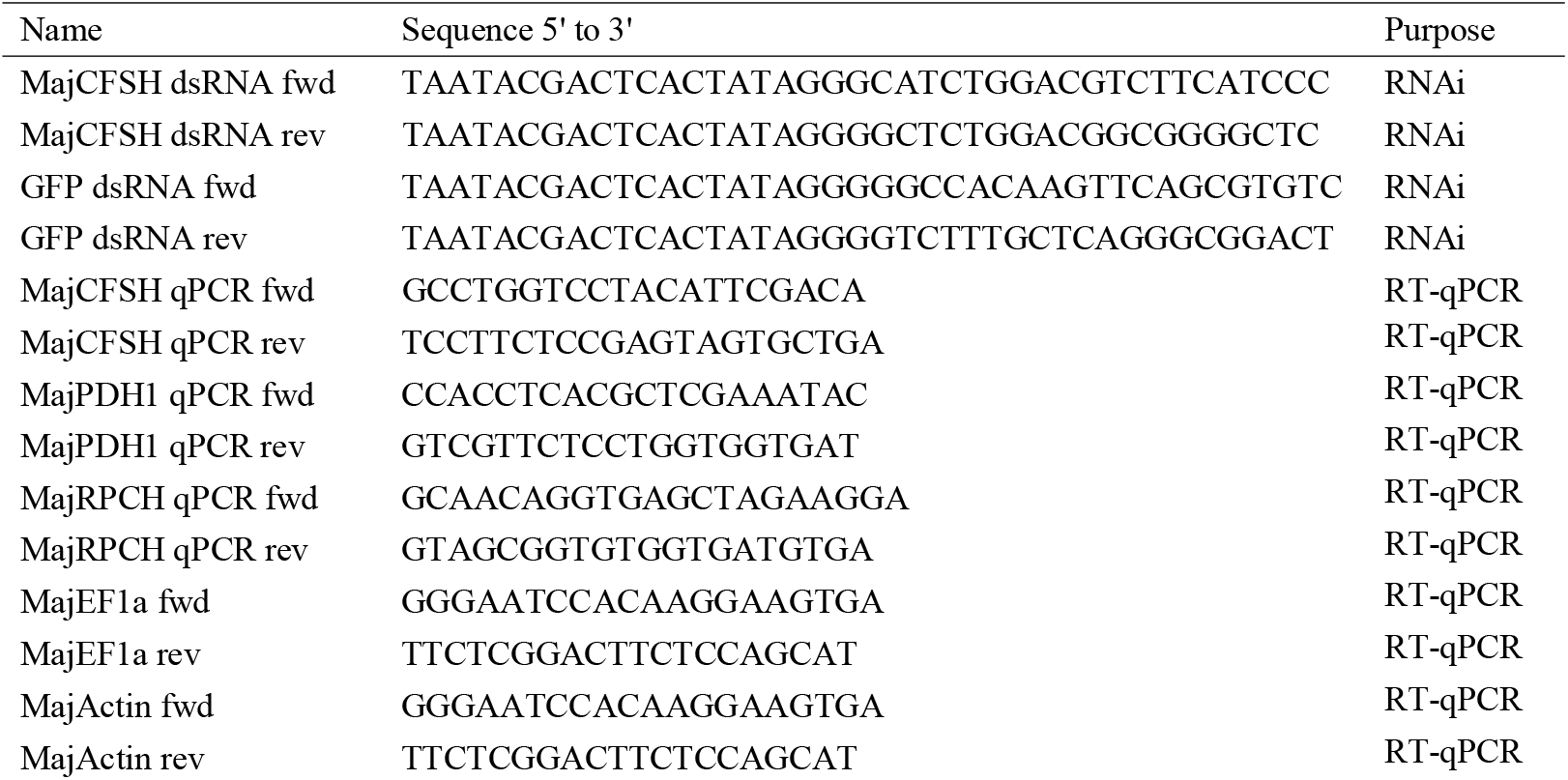
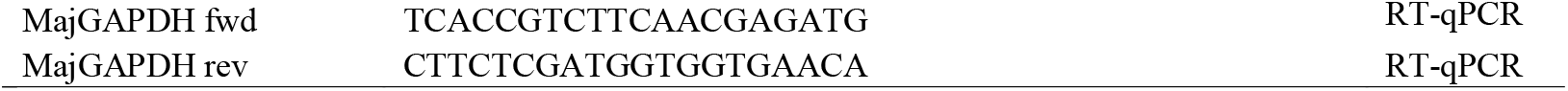
The primers used in this study.

### Quantitative PCR

Total RNAs of eyestalk samples from randomly selected juveniles (n = 12) of each group (GFP-dsRNA-injected female, GFP-dsRNA-injected male, CFSH-dsRNA-injected female, CFSH-dsRNA-injected male) were extracted and treated with DNase using the ISOGEN II (Nippon Gene, Tokyo, Japan) and RNeasy Micro Kit (Qiagen, Tokyo, Japan). The cDNA was synthesized from 250 ng of total RNA using SuperScript III with random hexamer primer (Life Technologie), according to the manufacturer’s protocol. RT-qPCR was conducted with TB Green Premix Ex Taq II using the ABI Prism 7000 thermocycler (Life Technologies). PCR conditions were as follows: 95°C for 10 min and 45 cycles at 95°C for 30 sec, 58°C for 1 min, and 72°C for 1 min in 20 μl volumes. The primer sequences used for the quantitative PCR analyses are shown in Table 1. The accession numbers of pigment dispersing hormone 1 (MajPDH1) and red pigment concentrating hormone (MajRPCH) were AB073369 and XP_042873530, respectively. Among three housekeeping genes (actin, GAPDH, EF1α), EF1α was selected as the reference gene calculated by NormFinder software (Andersen et al., 2004). Statistical analysis of gene expression was performed by Welch’s t-test using R (R Core Team, 2022).

## Results

### MajCFSH is involved in the growth of male juveniles

The RNAi experiment started with 80 juveniles in each injection group, but only 33 juveniles (41.3%) of the GFP-dsRNA group and 44 (55.0%) of the CFSH-dsRNA group survived to the end of the experiment (6 weeks). Identification of either females or males at the end of the experiment was conducted by external morphology of the spermatheca for females **(Figure S1B)** and the first pleopods for males **(Figure S1C)**. No individuals were difficult to distinguish as male or female by external morphology, such as having both male and female traits. There was no significant difference in survival between males and females in the two groups (GFP-dsRNA female =21, CFSH-dsRNA female =19, GFP-dsRNA male =12, CFSH-dsRNA male =25), suggesting that the fortnightly injections were responsible for the lower overall survival rate. The RNAi efficiency of *MajCFSH* was assessed by RT-qPCR using eyestalks from 7 females and 5 males of the GFP-dsRNA-injected group, and 6 females and 6 males of the MajCFSH-dsRNA-injected group, with the result that the expression of *MajCFSH* in both sexes was significantly suppressed in the MajCFSH-dsRNA group **(Figure 1)**. Males showed significantly lower body length and weight in the MajCFSH-dsRNA group compared to the GFP-dsRNA group, while no difference was observed in females in both groups **(Figure 2)**.

**Figure 1.**
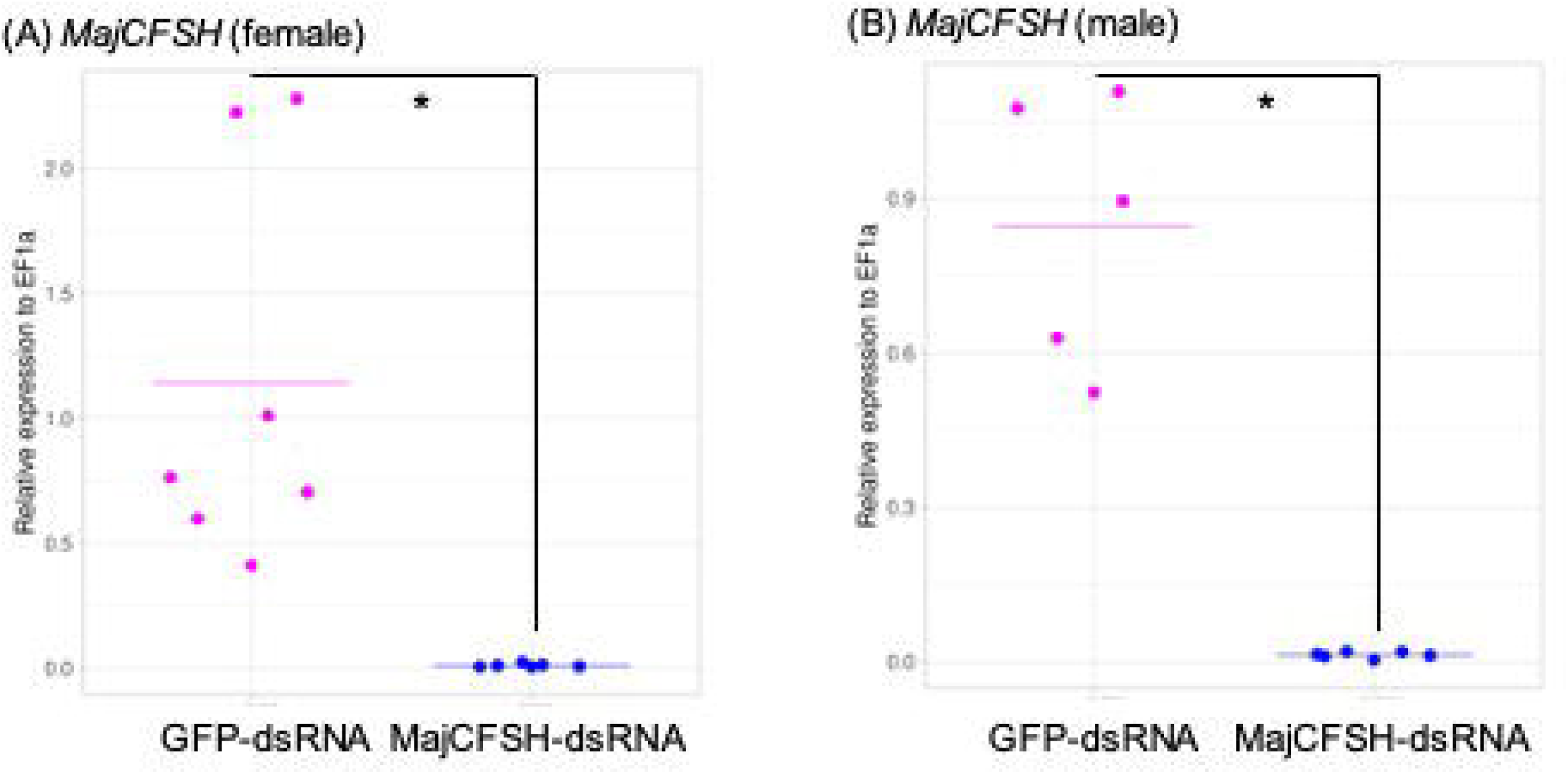
Expression profiles of *MajCFSH* (A, B) in the GFP- or CFSH-dsRNA-injected juveniles, respectively. The number of samples per condition is: GFP-dsRNA female =7, CFSH-dsRNA female =6, GFP-dsRNA male =5, CFSH-dsRNA male =6. Upper and lower showed the results of females and males, respectively. The asterisks denote significant differences between the GFP- and CFSH-dsRNA-injected juveniles (Welch’s t-test, p < 0.05).

**Figure 2.**
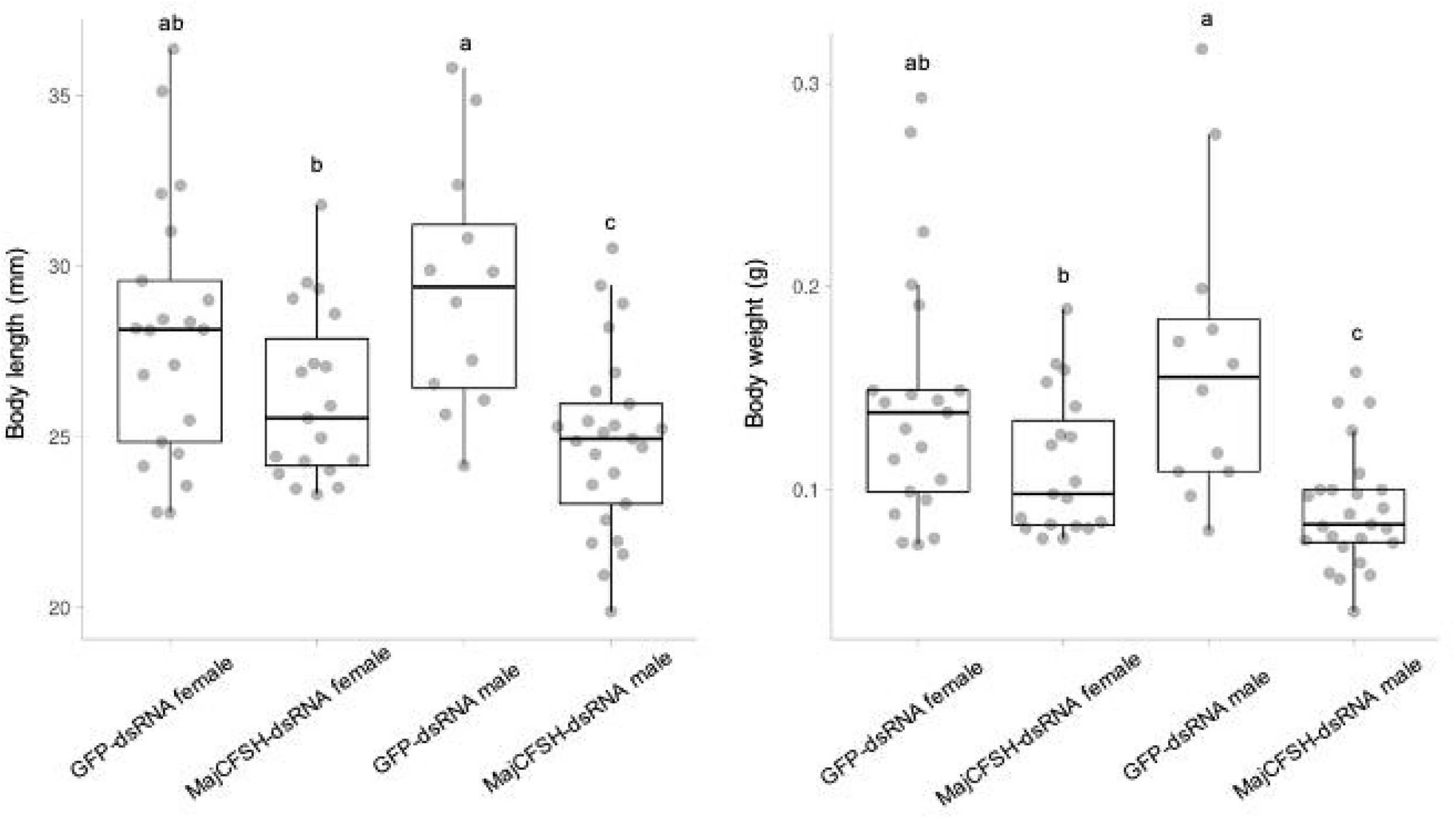
Average body length and weight at the end of RNAi experiment (six weeks) were indicated in (A) and (B), respectively. The number of samples per condition is: GFP-dsRNA female =21, CFSH-dsRNA female =19, GFP-dsRNA male =12, CFSH-dsRNA male =25. The different letters denote significant differences among four categories (One-way ANOVA, Post hoc Tukey–Kramer test, p < 0.05).

### MajCFSH has the potential to change the body color of juveniles

After the RNAi experiment, we noticed that the body color of the MajCFSH-dsRNA group was brighter/lighter **(Figures 3A, 3B, 4A, and 4B)**. Then, we extracted body color information from the dorsal parts **(Figure S1A)** and transformed it into HSV (Hue, Saturation, Value) color space **(Figures 3C and 4C)**. Comparison of H, S, and V, separately, between the two groups revealed significant changes in all three indices in both females **(Figures 3D-F)** and males **(Figures 4D-F)**.

**Figure 3.**
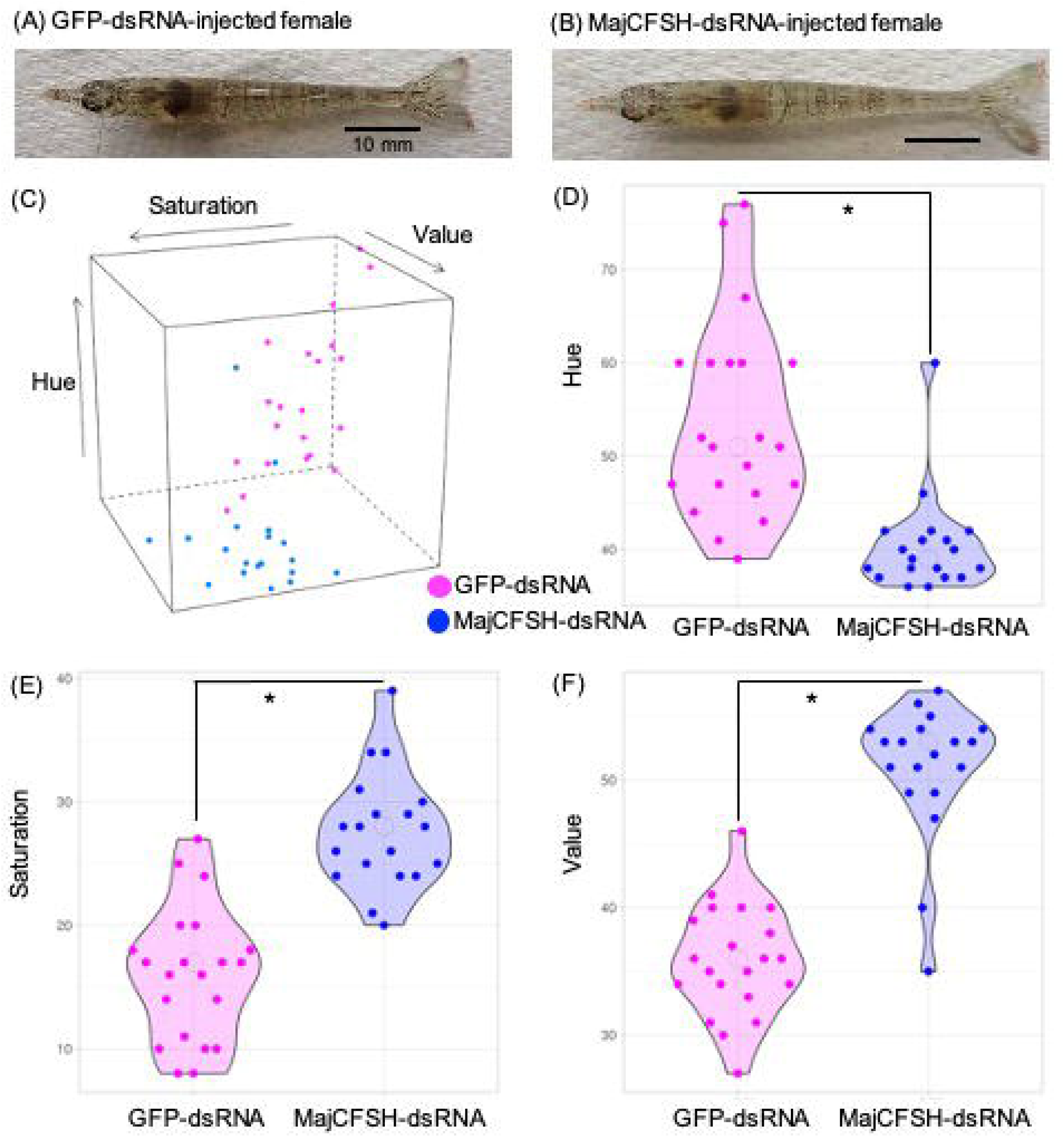
Body color change by CFSH-knockdown in female juveniles (GFP-dsRNA female =21, CFSH-dsRNA female =19). Snapshots of GFP- or CFSH-dsRNA-injected females just before sacrifice, respectively (A, B). Distribution of average body color in the HSV color space (C). HSV values of body colors were analyzed using a digital camera (TG-6, Olympus) and photoshop software. Hue, Saturation, and Value were shown in (D, E, F). respectively. *; P < 0.01 (Welch’s t-test).

**Figure 4.**
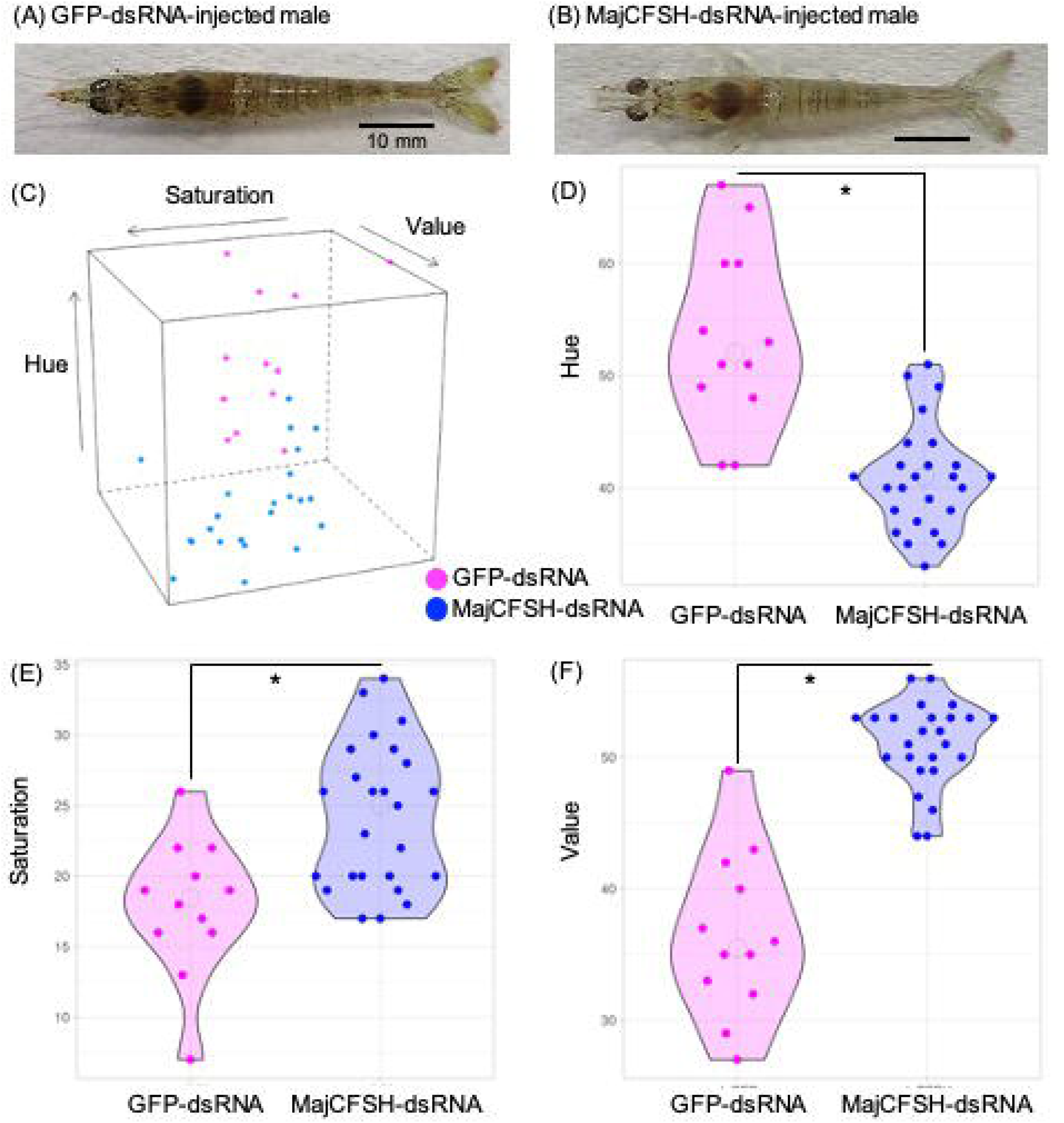
Body color change by CFSH-knockdown in male juveniles (GFP-dsRNA male =12, CFSH-dsRNA male =25). Snapshots of GFP- or CFSH-dsRNA-injected females just before sacrifice, respectively (A, B). Distribution of average body color in the HSV color space (C). HSV values of body colors were analyzed using a digital camera (TG-6, Olympus) and photoshop software. Hue, Saturation, and Value were shown in (D, E, F). respectively. *; P < 0.01 (Welch’s t-test).

### MajCFSH has the potential to regulate body color change in juveniles

To elucidate the mechanism of MajCFSH-RNAi-induced body color changes, the expression levels of other well-known color regulators, *pigment dispersing hormone 1* (*MajPDH1*) and *red-pigment concentrating hormone* (*MajRPCH*), were examined in the eyestalks of GFP- or MajCFSH-dsRNA-injected groups. The results showed a tendency for *MajPDH1* expression levels to increase in the CFSH-dsRNA-injected group in both sexes **(Figures 5A, 5B)**, while no change in *MajRPCH* was observed in either sex in each injected group **(Figures 5C, 5D)**.

**Figure 5.**
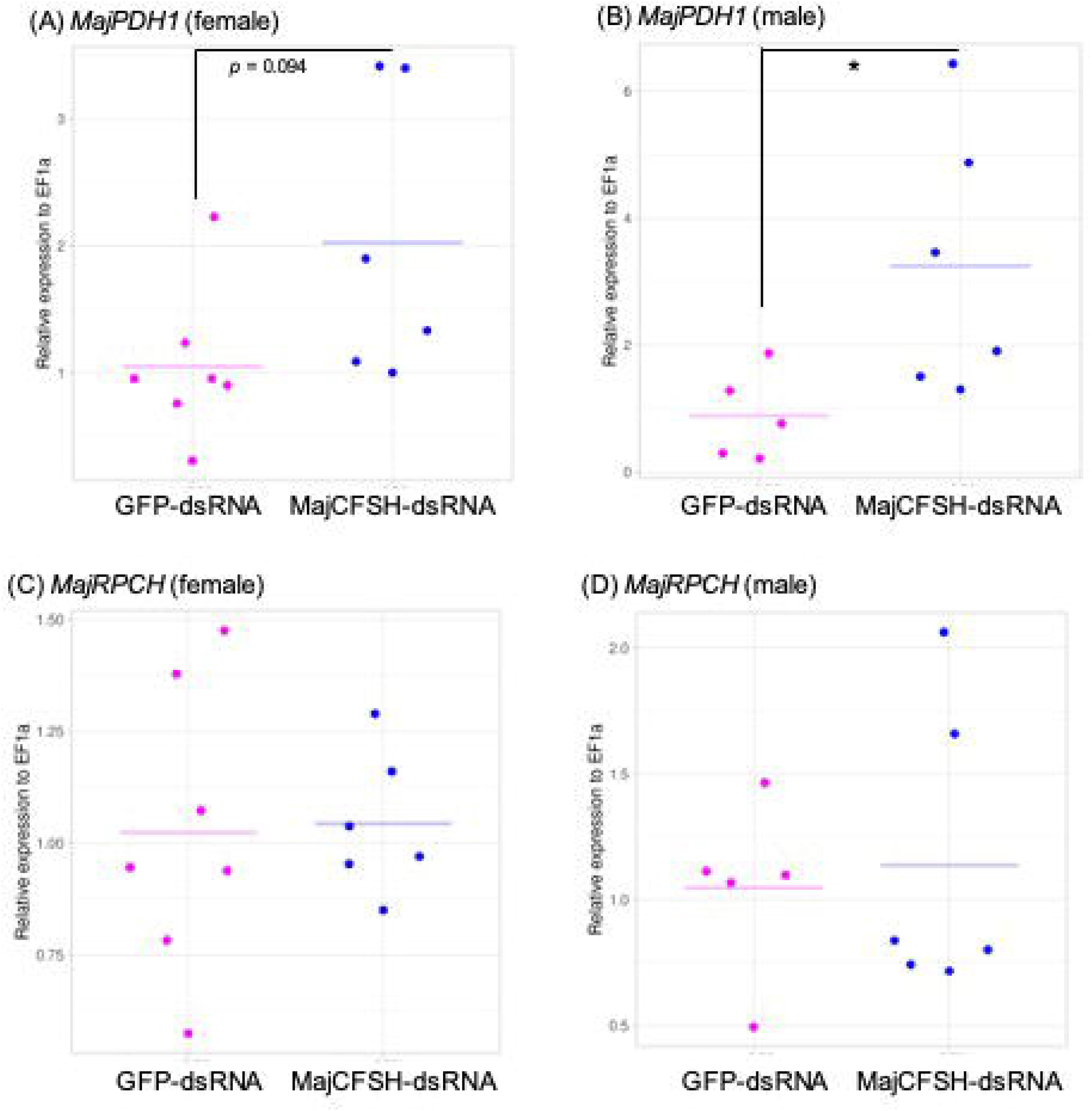
Expression profiles of *MajPDH* (A, B) and *MajRPCH* (C, D) in the GFP- or CFSH-dsRNA-injected juveniles, respectively. The number of samples per condition is: GFP-dsRNA female =7, CFSH-dsRNA female =6, GFP-dsRNA male =5, CFSH-dsRNA male =6. Upper and lower showed the results of females and males, respectively. The asterisks denote significant differences between the GFP- and CFSH-dsRNA-injected juveniles (Welch’s t-test, p < 0.05).

## Discussion

The present study demonstrated that eyestalk-type MajCFSH has dual functions; one is male-specific growth regulation, and another is body color regulation in both female and male juveniles. The growth of crustaceans is centrally regulated by molting because their cuticle-based hard shell inhibits continuous growth, such as in vertebrates. Molting of Malacostracan crustaceans is regulated by molt-inhibiting hormone (MIH), which is a neuropeptide hormone secreted from the X-organ/sinus gland (XO/SG) complex in the eyestalks. MIH normally suppresses the biosynthesis of molting hormone (20-hydroxyecdysone; 20E), while before molting MIH expression is decreased dramatically, and then 20E biosynthesis and secretion take place, resulting in molting. Based on this molting regulation by MIH, eyestalk ablation is a widely accepted way to promote the growth in decapod species (Qiao et al., 2018), however, no XO/SG-derived neuropeptides, except for MIH, have been reported to regulate juvenile growth. In this study, MajCFSH-knockdown males obviously showed growth suppression, suggesting that MajCFSH might contribute to male-specific juvenile growth or/and trait development. Female kuruma prawns have a higher fisheries value than males because females grow faster. However, if MajCFSH can promote male juvenile growth, it may be possible to increase the fisheries value of male kuruma prawns. As a next step, an injection experiment of purified or recombinant MajCFSH into kuruma prawn juveniles will be required.

Research into the role of CFSH in male sexual differentiation is still in the early stages because it has been identified as a female-specific hormone that plays a crucial role in female phenotype development in crustaceans (Zmora and Chung, 2014; Toyota et al., 2021). Recently, several works have demonstrated the interaction of CFSH and other factors involved in sexual differentiation. Insulin-like androgenic gland factor (IAG) is a primary molecule for male sexual differentiation and the development of secondary male sex characteristics and is synthesized and secreted from the androgenic gland (AG), which is a male-specific endocrine organ (Levy and Sagi, 2020). AG transplantation or promotion of IAG expression induces masculinization, while AG removal or suppression of IAG expression results in partial or complete functional sex reversal in several decapods (Katayama et al., 2022; Kato et al., 2015; Sagi et al., 1990; Rosen et al., 2010; Ventura et al., 2011). A few studies have suggested the presence of an endocrine axis connecting eyestalk ganglia and AG in male decapods, because eyestalk ablation triggers rapid hypertrophy of the AG and up-regulation of IAG expression in *C. sapidus* (Chung et al., 2011) and the red-claw crayfish *Cherax quadricarinatus* (Khalaila et al., 2002). In some decapod research, a CFSH inhibitory effect has been demonstrated on AG activity that synthesizes and secretes IAG (Jiang et al., 2020; Liu et al., 2018; Tsutsui et al., 2018), suggesting that CFSH might be one of the eyestalk-derived neuropeptides that are involved in male sexual differentiation via modulating IAG expression in the AG.

More recent sex differentiation studies of decapods have also focused on the role of vertebrate sex steroid hormones in addition to CFSH and IAG. Indeed, 17β-Estradiol (E2) and testosterone (T) have been detected in *C. sapidus* hemolymph by ELISA and LC-MS/MS methods, and several homologous genes involved in their biosynthesis have been identified (Wang et al., 2022, 2023). In addition, regulatory relationships among CFSH, IAG, E2, and T are gradually becoming clear from knockdown experiments of CFSH and IAG. The effects of E2 and T on sex differentiation will be verified in the kuruma prawn, and the substantial network between CFSH and other sex differentiation factors (IAG, E2, and T) will also be clarified in the future. Furthermore, the physiological function of ovary-type MajCFSH-ov, which has been identified in the kuruma prawn, needs to be clarified using the RNAi approach.

Decapod chromatophores are pigment-containing cells found in the epidermis of decapod crustaceans such as shrimp and crabs (Elofsson and Kauri, 2022). These cells are responsible for the dynamic optical response of biological tissue, allowing them to camouflage, shade, and thermoregulate (Kay et al., 2022). Chromatophores contain pigment granules that can be moved around within the cell by a process known as pigment aggregation (Ribeiro and McNamara, 2007). This process is regulated by neurosecretory peptides and involves an increase in intracellular calcium (Ribeiro and McNamara, 2007). Red pigment-concentrating hormone (RPCH) and pigment-dispersing hormone (PDH) are crustacean neuropeptides that are involved in a variety of physiological processes, including body color changes in decapods (Ohira et al., 2002; Wei et al., 2021). RPCH influences the concentration of pigment chromatophores, causing drastic body color changes in crustacean species. Studies have shown that changing body color in decapods into paler colors is influenced by rearing conditions such as light intensity and temperature (Ranga et al., 2001). In this study, *MajCFSH* knockdown induced body color changes to become brighter than GFP-dsRNA-injected juveniles, indicating that MajCFSH has the potential to disperse the pigment granules in the chromatophore, like PDH. The trend toward increased *MajPDH1* expression in the eyestalks of *MajCFSH*-knockdown individuals was thought to restore the lightened body color to its original state. Either way, body color lightening was caused by *MajCFSH* knockdown, but no decrease in *MajPDH1* expression nor any increase in *MajRPCH* expression was observed. Based on these results, it can be interpreted that MajCFSH does not change body color via MajRPCH or MajPDH1, but that MajCFSH itself has the ability to change body color. Therefore, MajCFSH is a novel body color-regulating hormone in decapod crustaceans other than RPCH and PDH.

## Conclusion

This study investigated the physiological roles of MajCFSH by RNAi experiment. Although it is not involved in the development of sex-specific characteristics such as external reproductive organs, MajCFSH-knockdown male juveniles showed apparent growth suppression, implying that MajCFSH may play a male-specific juvenile growth role. Moreover, MajCFSH-knockdown female and male juveniles changed their body color to become brighter, indicating that MajCFSH has the ability to change body color by dispersing the pigment granules in the chromatophore. Overall, our present study improved our understanding of the physiological roles of CFSH in the kuruma prawn.

## Declaration of Competing Interest

The authors have no competing interest to declare.

## Author contributions

KT and TO designed the conception of this study. All authors conducted the experiment, material preparation, data collection, and discussion. KT analyzed the all data for publication. The first draft of the manuscript was written by KT and all authors commented on it. All authors read and approved the final manuscript.

## Acknowledgments

The authors would like to thank Dr. Mike Roberts (Independent Consultant, UK) for his critical readings of this manuscript. This study was supported in part by grants to TO (Grant-in-Aid for Scientific Research [B] No. 22H02439 by JSPS), and the Japan Science and Technology Agency/Japan International Cooperation Agency, Science and Technology Research Partnership for Sustainable Development, SATREPS (JPMJSA1806), Project for Utilization of Thailand Local Genetic Resources to Develop Novel Farmed Fish for Global Market (Thai Fish Project).

## Supplemental data

Figure S1

P1 to P3 indicate the sites for which HSV values were obtained (A). Abdominal morphology of females (B) and males (C). A’ and B’ show the spermatheca and the first pleopod with white-dotted outlines, respectively.

## Notes

### Competing Interest Statement

The authors have declared no competing interest.

