## Supplementary figures and images for "Dual roles of crustacean female sex hormone during juvenile stage in the kuruma prawn *Marsupenaeus japonicus*"

### FigS1.jpg

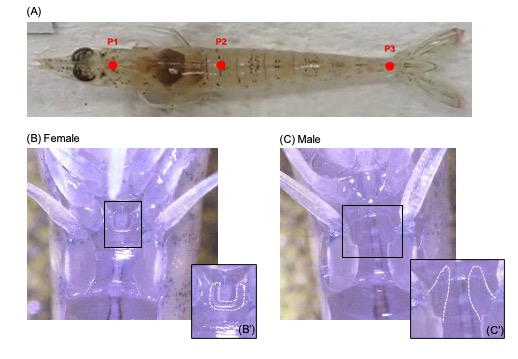
